# Mosquito virus diversity in Western and North-western Uganda

**DOI:** 10.1101/2023.11.02.565349

**Authors:** Martin N. Mayanja, Marc Niebel, Shirin Ashraf, Maryam Hardy, Alfred Ssekagiri, Lily Tong, Anne Nanteza, Enock Matovu, Sylvester Nyakaana, Ana da Silva Filipe, Frank Mwiine, Julius J. Lutwama, Emma C. Thomson

## Abstract

**Introduction:** Surveillance for mosquito borne arboviruses in Sub Saharan Africa has largely focussed on known viruses including Yellow fever virus (YFV), Rift Valley fever virus (RVFV), Chikungunya virus (CHIKV), West Nile virus (WNV) and Dengue viruses (DENV). Routine surveillance and outbreak investigations traditionally rely on serology, PCR and cell culture. Although such methods are useful, they are do not detect novel or unexpected viruses.

**Methods:** This study employed unbiased metagenomic next generation sequencing (MNGS) to characterise viruses circulating in mosquitoes of Arua and Kasese districts of Uganda collected systematically as part the ArboViral Infection (AVI) study. Adult mosquito sampling was carried out from multiple sites using light traps baited with solid carbon dioxide (indoors) and pyrethrum spray (outdoors). 10,026 mosquitoes were identified using appropriate morphological identification keys and separated into 96 pools by species and location of collection. Viral RNA extracted from homogenised mosquitoes was reverse transcribed to complimentary DNA and sequenced on the flow cell of an Illumina NextSeq platform. Bioinformatic analysis was performed using a customized in-house metagenomics pipeline. Downstream analyses were carried out in R version 4.2.1.

**Results:** 97 viruses from 24 families and 31 genera were detected in 96 mosquito pools from Arua and Kasese districts. The abundance of viruses in different families was in the order *Rhabdoviridae* (33), *Flaviviridae* (28), *Orthomyxoviridae* (13), *Mesoniviridae* (9), *Piconarviridae* (9), *Peribunyaviridae* (9), *Phasmaviridae* (9), *Iflaviridae* (8), *Phenuiviridae* (7), *Nodaviridae* (5), *Xinmoviridae* (4), *Reoviridae* (4), *Virusidae* (2), *Nairoviridae* (2), *Qinviridae* (2), *Togaviridae* (2), *Alphatetraviridae* (2), *Picornaviralesidae* (2), *Iridoviridae*, (1), *Nudiviridae* (1), *Parvoviridae* (1), *Permutetraviridae* (1) and 22 viruses from different families remain unclassified. The *Flaviviridae*, *Togaviridae*, *Rhabdoviridae*, *Peribunyaviridae*, *Phenuiviridae*, *Nairoviridae, Nodaviridae* and *Orthomyxoviridae* harbor viruses that cause disease in humans and other mammals. Viruses from insect-specific virus families that were detected included the *Chuviridae*, *Iflaviridae*, *Phasmaviridae and Alphatetraviridae*. 92/173 (53%) of the viral genomes had full opening reading frames (ORFs) and diverged by >30% nucleotide pairwise distance from the nearest reference genome.

**Conclusion:** The majority of viruses detected were novel species described for the first time with unknown potential to cause disease. The diversity of virus species in mosquitoes from Uganda, a hotspot for emerging arboviruses has been only partially characterized. This study, carried out in Western and North West Uganda illustrates the scale of richness and virus diversity in the region and the need to further characterise the virome in mosquitoes, especially those with a propensity to feed from human and animal hosts.

## Introduction

Despite the well-documented richness of viral species diversity in Uganda, there have to date been few studies employing agnostic methods to characterise the virome of mosquitoes in this region (Marklewitz et al., 2013; Sempala, 2002). Currently, 24 mosquito-borne viruses affecting different animal hosts classified in several families have been documented in Uganda (Mayanja et al., 2021). The majority have been identified using traditional cell-culture, PCR or serological methods. Although such experimental approaches have been successful in documenting a proportion of viral species, they have important limitations. Some viruses may be challenging to culture, and PCR-based methods rely on prior knowledge of the target nucleotide sequences (Kayiwa et al., 2018; Steinhagen et al., 2016). Unbiased metagenomic next generation sequencing (MNGS), now a well-established technique, has been used in the discovery and tracking of viruses from different animal host and vector species (Marklewitz et al., 2013; Ramesh et al., 2019), including in Uganda (McMullan et al., 2012). In this investigation, carried out as part of the ArboViral Infection study, we employed unbiased next generation sequencing to detect known and previously uncharacterised viruses in mosquitoes collected from multiple locations in Western and North Western Uganda.

## Materials and Methods

### Sample collection

Mosquito collections were carried out in Arua and Kasese districts in North Western and Western Uganda respectively and mosquito species were identified, as previously described (Mayanja, Mwiine, Kohl, Thomson, & Lutwama, 2020). In Arua district, sample collection was carried out in the four villages of Yedu, Oniba, Ambala and Barize while in Kasese district, it was done in Kidodo, Kirembe, and Kyondo and Road barrier villages.

### Viral RNA extraction and NGS

Mosquitoes were pooled by sex, species and location (**Supplementary Table 1**). Pools were homogenized using steel beads (5mm) in 1.3 ml of BA1 medium (5% Bovine Serum Albumin, 10X M199 Hanks salts, 1M Tris HCl, 7.5% Sodium bicarbonate, 100X Penicillin and Streptomycin, 1 ml of fungizone) using a gene grinder. The supernatant was clarified in a micro-centrifuge at 8000 rpm for 15 minutes and stored at −80C.

RNA was extracted from 400 µl sample using the Agencourt RNAdvance Blood extraction kit (Beckman Coulter) following the manufacturer’s protocol including a 15 min DNase treatment at 37C (Ambion, Austin, TX, USA) and eluted in 20 µl of nuclease free water. 15 µl of extracted RNA was reverse transcribed using Superscript III enzyme (Invitrogen) and random hexamers. The product was converted to double stranded DNA using the NEBNext second strand synthesis kit (New England Biolabs). Library preparation was carried out using a KAPA LTP kit (KAPA Biosystems). Indexed libraries were quantified by Qubit fluorometry and sized on the Tape station 4200 (Agilent Technologies) and pooled in equimolar concentrations for sequencing on the Illumina NextSeq.

#### Bioinformatics and statistical analysis

NGS data analysis was performed on computational infrastructure provided by the MRC-University of Glasgow Center for Virus Research (CVR). Raw fastq file reads were inspected using DIAMOND BLASTx, coupled with Krona tools in a custom script (Buchfink, Xie, & Huson, 2015). Krona plots were visually inspected using Firefox browser and later translated to text output files showing the taxonomy of viruses detected by Diamond BLASTx. Complete reference genomes of known viruses of interest detected during exploratory analysis were downloaded from NCBI. Raw fastq files were mapped to each of the reference sequences using Tanoti version 1.3 (https://github.com/vbsreenu/Tanoti). Mapping statistics and consensus sequences were generated from sequence alignment maps using samtools. Results of reference-based mapping were interrogated by examining the number of mapped reads and percentage coverage using the ggplot2 R package. *De novo* assembly was carried out using the Metavic pipeline (https://github.com/sejmodha/MetaViC) which incorporates SPAdes, IDBA-UD for trimming, assembly and merging of paired reads and GARM for generating supercontigs. Contigs from *Denovo* assembly were blasted against RefSeq protein from NCBI using DIAMOND BLASTx. Html files were visualized using Firefox browser and then translated into text output files. Consensus sequences identified as viral were identified by BLASTn. Tanoti was used for reference-based mapping to novel contigs generated by *de novo* assembly.

Using IQtree, a supertree for the different viruses from different families was created. An Evolutionary Phylogenetic Algorithm PA-NG (EPA-NG) was then used to place the multiple set of non-overlapping contigs onto the underlying phylogeny as shown (Figure 1). One contig per mosquito pool per virus was used in order to represent the distribution of different species in mosquitoes. Visualization was implemented using Interactive Tree of Life (iTOL).

**Figure 1:**
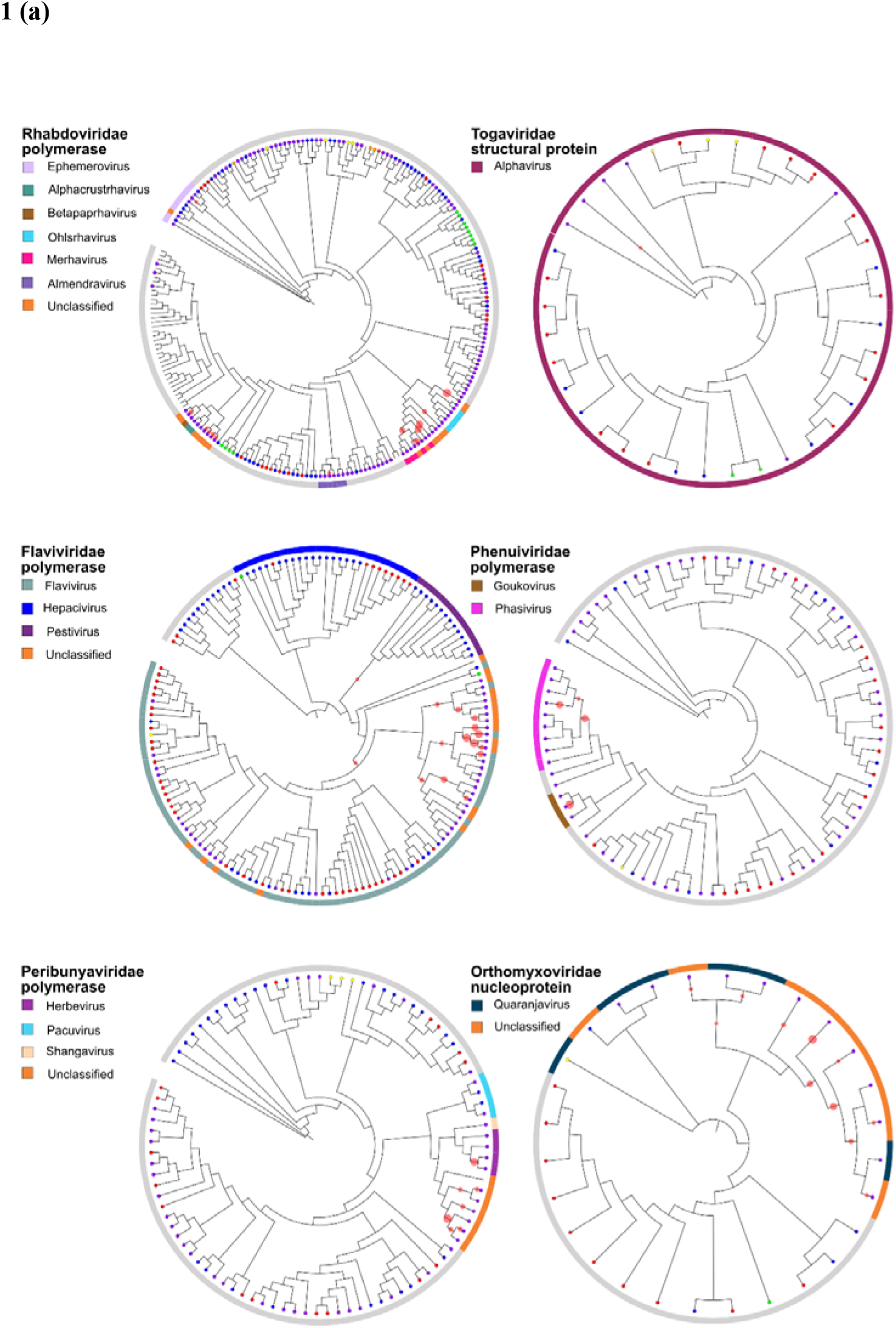
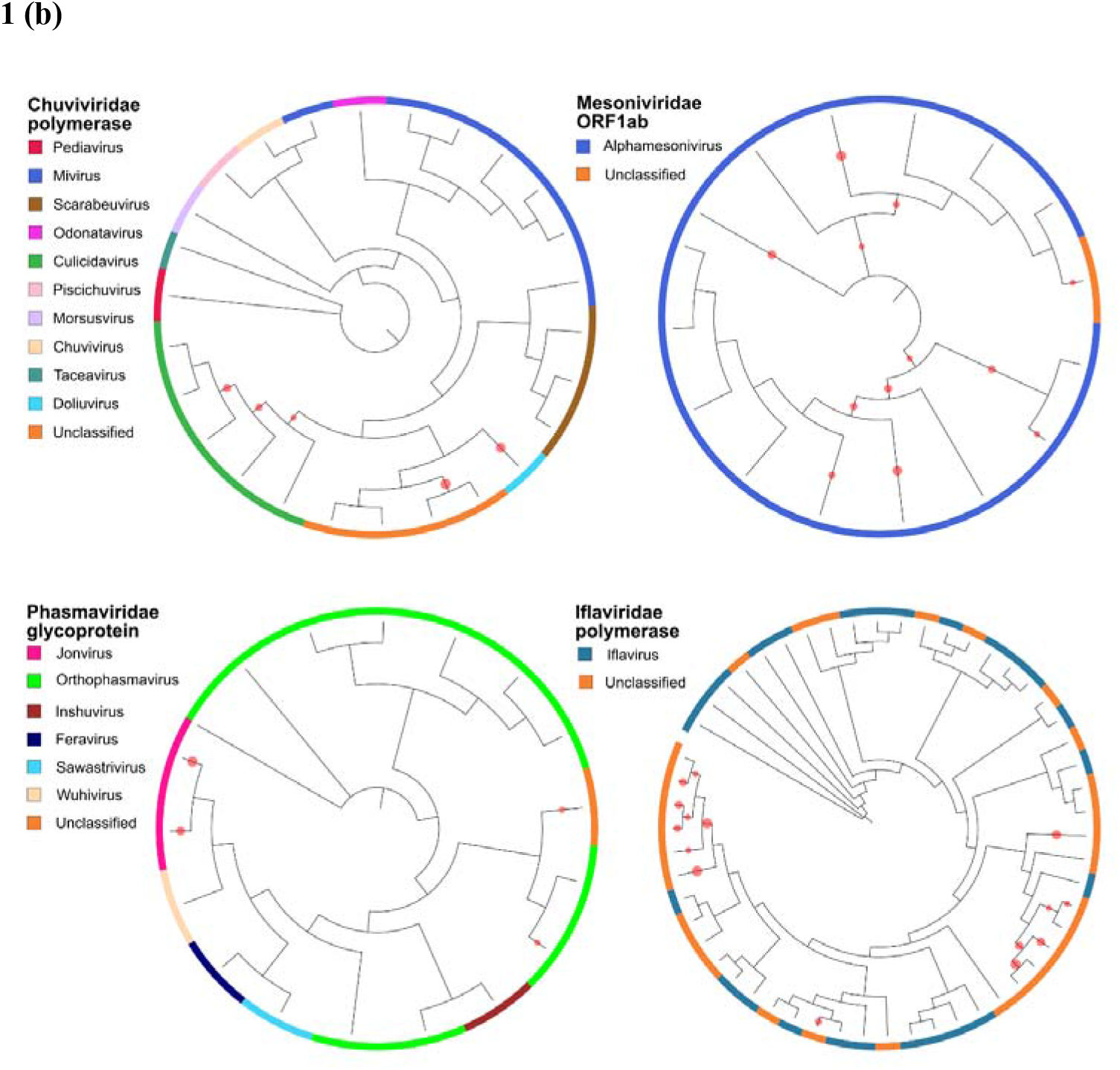
Virus diversity in mosquitoes from Western and North Western Uganda. Virus sequences with homology to viruses in families with species known to infect (a) **Human and animal hosts** and (b) **Restricted to arthropod hosts**

Multiple sequence alignment was carried out using MAFFT. Maximum likelihood phylogenetic trees were drawn for near complete amino acid genomes using IQTREE and RaxML. Uncorrected nucleotide and amino acid pairwise distances were calculated on MAFFT alignments using MEGA-X. Criteria for novel viruses were taken from ICTV and/or relevant literature and were restricted to viruses with a contig length of >1kb.

Virus genus diversity was summarized using heat maps. Counts of virus hits were aggregated by genus and family of the closest reference; distribution of virus genera and families were visualized using the pheatmap package. The *estimate-richness* function of the phyloseq package was used to compute alpha diversity of virus genera across sampling locations and mosquito genera. The distribution of the richness and Simpson’s index was compared amongst sampling locations and mosquito genera using Kruskal Wallis test. Multiple pairwise comparisons of alpha diversity amongst mosquito genera were done using Dunn’s test. We used the ggplot2 package for visualization. Analysis was carried out in R computing environment version 4.2.1.

## Results

A total of 96 mosquito cDNA pools were included in the study and an average of 5,978,259 million sequence reads was generated per pool (**Supplementary Table 1**). The length of contigs mapping to viruses ranged from 230 to 21,147bp. Extensive, previously undocumented viral diversity and high virus richness was noted in all mosquito species sampled. There were no differences between virus genera Simpson’s diversity index amongst mosquito genera (adjusted pairwise comparison p-values). The virus diversity and richness was also not different between areas sampled. A total of 173 virus RNA sequences from 25 families and 42 genera were detected. Out of these, 92 (53%) diverged by 30% nucleotide pairwise distances, suggestive of novel viral species (**Supplementary Table 2**).

The abundance of viruses in different families was in the order *Rhabdoviridae* (33), *Flaviviridae* (28), *Orthomyxoviridae* (13), *Mesoniviridae* (9), *Piconarviridae* (9), *Peribunyaviridae* (9), *Phasmaviridae* (9), *Iflaviridae* (8), *Phenuiviridae* (7), *Nodaviridae* (5), *Xinmoviridae* (4), *Reoviridae* (4), *Qinviridae* (2), *Virusidae* (2), *Nairoviridae* (2), *Togaviridae* (2), *Alphatetraviridae* (2), *Picornaviralesidae* (2), *Iridoviridae*, (1), *Nudiviridae* (1), *Parvoviridae* (1), *Permutetraviridae* (1) and 22 unclassified viruses from different families. Viruses were identified in a total of 44 virus genera. Nine virus genera were unique to Kasese (*Ledantevirus, Orthobunyavirus, Lentivirus, Pegivirus, Orbivirus, Perhabdovirus, Almendravirus, Vesiculovirus* and *Chloridovirus*. Similarly, six virus genera were unique to Arua (*Enterovirus, Lyssavirus, Dinovernavirus, Sigmavirus, Alphanemrhavirus, Aquabinavirus*).

In order to illustrate the spread within virus families of non-overlapping contigs detected in this study, we used a phylogenetic placement program (EPA-NG) following alignment of contigs to pre-existing amino acid trees derived from references sequences for each family. The longest contig from each virus was selected for every sample, to facilitate a quantitative representation of viruses within each mosquito pool (**Figure 1**). Among the 25 virus families, the *Flaviviridae*, *Togaviridae*, *Rhabdoviridae*, *Peribunyaviridae*, *Phenuiviridae* and *Orthomyxoviridae* are known to harbor human and animal pathogens (**Figure 1a**). Other families detected included the arthropod-specific virus families *Chuviridae*, *Iflaviridae*, *Phasmaviridae* and *Mesoniviridae* (**Figure 1b**). The EPA-NG algorithm was used to place non-overlapping contigs for different virus families onto a protein phylogenetic tree in order to illustrate the diversity of viruses detected. This does not allow identification of novel viruses, but provides a visual illustration of within-family diversity.

In order to identify new and existing species, maximum likelihood phylogenetic trees were created (**Figure 2**). Full-length open reading frames of viruses from several viruses were detected, several of which reached the criteria for novel viruses.

**Figure 2:**
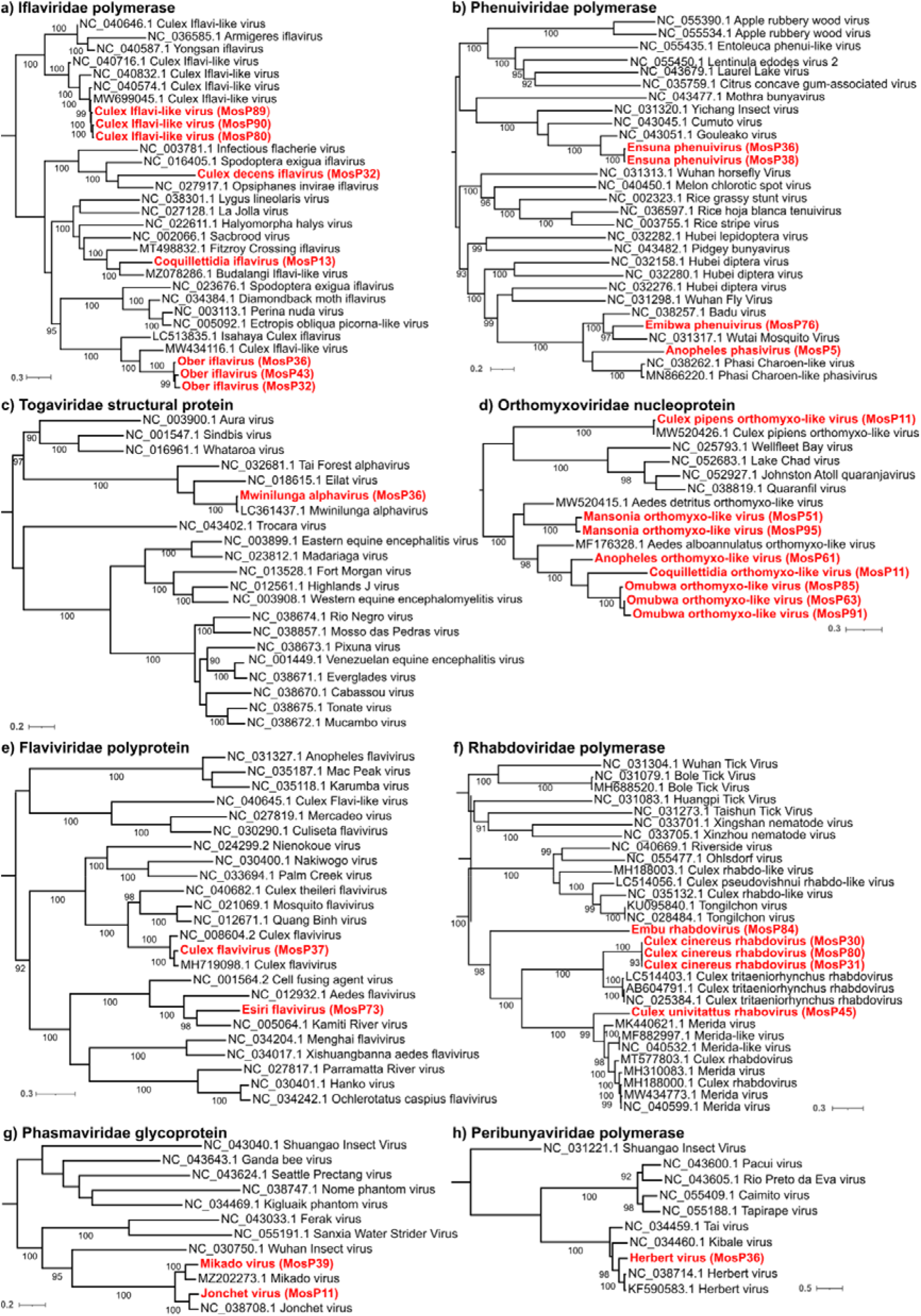
Maximum likelihood phylogenetic trees of full open reading frames detected within key virus families

### *Togaviridae* family

Family *Togaviridae* consists of positive-sense single stranded RNA viruses vectored infecting a wide range of animal hosts. In Uganda, five different viruses have been documented. Mwinilunga alphavirus (LC 361437.1) that was first identified in *Culex quinquefasciatus* from Zambia was also detected in pool MosP36 of *Culex duttoni* collected from Arua (Torii, 2018 #562). This is the first report of Mwinilunga alphavirus in Uganda (Figure 2c). The query virus in pool 36 had equal branch length with MWAV, a closely related to Eilat virus mosquito-specific alphavirus.

### *Orthomyxoviridae* family

Family *Orthomyxoviridae* consists of RNA segmented (6 to 8 segments) viruses categorized in seven genera. Viruses in this family have got a worldwide distribution infecting animals, humans and birds. Out of the seven genera, genera Quaranjavirus and Thogotovirus are arboviruses transmitted either by mosquitoes or ticks. This is the first report of an orthomyxovirus detected in mosquitoes in Uganda. A total of five orthomyxoviruses, four of which were novel viruses, that have been named Anopheles orthomyxo-like virus, Coquillettidia orthomyxo-like virus and Mansonia orthomyxo-like and Omubwa orthomyxo-like viruses were detected in pool MosP 61 of Anopheles spp, pool MosP 11 of Coquillettidia spp, a pool of Anopheles and Culex spp and pools MosP51, MosP95 of Mansonia uniformis and Omubwa orthomyxo-like virus from pools MosP 63, MosP85 and MosP91 of Anopheles and Culex mosquitoes respectively. Phylogenetic analysis of all the novel viruses with reference orthomyxovirus genomes showed a close relationship with Aedes alboannulatus orthomyxo- like viruses.

Another virus Culex pipiens orthomyxo-like virus which clustered closely with Culex pipiens orthomyxo-like virus (MW520426.1) first detected in a pool of Culex pipiens was detected in pool MosP11 of Coquillettidia spp.

### Flaviviridae family

Flaviviridae family consist of positive sense single stranded RNA viruses categorized in five genera. Majority of these are vectored by arthropods including mosquitoes and ticks transmitted to animals as well as humans while others are insect specific viruses. To date, over 20 viruses in family Flaviviridae have been documented in Uganda. Phylogenetic analysis of query samples with other flaviviruses showed pool MosP37 of *Culex duttoni* with highest identity to Culex flavivirus reference strain (MH719098.1) that was identified in *Culex quinquefasciatus* from Mexico. A novel virus named Esiri flavivirus was identified in pool MosP73 of Culex spp from Kasese district. Culex flavivirus in pool 37 clustered and had equal branch length with Culex flavivirus (MH719098.1) that was identified in Culex quinquefasciatus from Mexico. Virus contig in mosquito pool 73, had a longer branch length and clustered with Kamiti river virus (NC 005064.1) an insect specific flavivirus first described in *Aedes mcintoshi* collected from the Central Province of Kenya (Crabtree, 2003 #561). Pairwise distance between the query virus and Kamiti river virus was 31.51% bp. Genome length of the newly described virus was 10917bp compared to the reference virus genome which is 11375bp.

### *Rhabdoviridae* family

Family *Rhabdoviridae* is comprised of negative sense single stranded RNA viruses infecting a wide range of animal hosts including plants. Viruses in this family are very diverse and categorised into 46 genera while others remain unclassified. In Uganda, only two Rhabdoviruses of Kamese virus and Yata virus have been detected in mosquitoes. Three novel rhabdoviruses we have named Embu rhabdovirus, Culex cinereus rhabdovirus and Culex univitattus rhabdovirus were identified in mosquitoes from Uganda. Amino acid phylogenetic tree for Rhabdoviruses based on the L protein sequences show virus sequences in pools MosP30, MosP31 and MosP80 of *Culex cinereus* form a separate group that clusters with reference strains of Culex tritaeniorhynchus rhabdovirus (LC514403.1, AB604791.1 and NC_025384.1) of genus *Merhavirus* first detected in Japan. Pairwise distance between Culex cinereus rhabdovirus and Culex tritaeniorhynchus rhabdovirus was, 30%. Pool MosP45 of Culex univitattus contained a novel virus we have named Culex univitattus rhabdovirus, this virus (11793 bp long) clustered with reference genome of Merida virus (MK440621.1) that was detected in Culex spp from Sweden. Pool MosP84 of Culex neavei from Kasese contained a rhabdovirus we have named Embu rhabdovirus (9867 bp long). Pairwise distance between query virus and reference virus strain was more than 30% suggesting novelty. This is the first report of Merhaviruses including an unclassified virus species in Africa later on Uganda.

### *Phasmaviridae* family

Family *Phasmaviridae* of Order Bunyavirales, comprises of six virus genera that infect exclusively arthropods. They are made of 3 segments (L, M and S segments) that are about 2000 to 7000 bp long. During this study, two viruses Mikado and Jonchet virus of genus *Jonvirus* were detected in Culicine mosquitoes. Amino acid phylogenetic tree based on the M segment revealed Mikado virus (MZ202273.1) in pool MosP39 of *Culex neavei* a virus that was first described in *Culex annulioris* in 2004 in in Cote d’Ivore. Another virus documented for the first time in Uganda was Jonchet virus (NC 038708.1) in pool MosP11 of *Coquillettidia* spp, a virus also first detected in Cote d’Ivore during the same study in 2004.

### *Peribunyaviridae* family

Family Peribunyaviridae consist of segmented virus categorized into… genera. Pool MosP36 of *Culex duttoni* contained Herbert virus a related virus which was first identified in Culex nebulosus in Cote d’Ivore.

### Iflaviridae family

Phylogenetic analysis showed three pools (MosP80, MosP89, MosP90) of *Culex cinereus*, *Culex pipiens* and *Culex pruina from Kasese district* contained Culex Iflavi-like virus 4 (MW699045.1) that had been detected in *Culex pipiens* in Belgium in 2019 (Wang, 2021 #556). Pool 32 contained novel iflaviruses, one that named Culex decens iflavirus closely related with Opsiphanes invirae iflavirus (NC_027917.1) and another virus that has been named Ober iflavirus that also was detected in pools MosP36 and MosP43 of *Culex decens* and *Culex univitattus* from Arua district.

We additionally detected three novel viruses we named s Culex decens iflavirus detected in *Culex decens* in Arua (pool MosP32), Coquillettidia iflavirus virus detected in *Coquillettidia fraseri* (pool MosP13) and Ober iflavirus detected in *Culex univitattus* and *Culex decens* of (MosP32 and MosP43), all in Arua district. Phylogenetic analysis between Coquillettidia iflavirus with other iflaviruses showed a close relationship with Fitzroy Crossing iflavirus 1 (MT498832.1). Pairwise distance between Coquillettidia iflavirus and Fitzroy Crossing iflavirus 1 first detected in Culex annulirostris from Western Australia was 41.67% (Williams, 2020 #338). Another novel virus Culex decens iflavirus detected in *Culex decens* in Arua (pool MosP32) clustered with Culex iflavi-like virus 3 first detected from Culex species in USA in 2016. (Sadeghi, 2018 #555). Pairwise distance between the two viruses was 45.46%.

### *Phenuiviridae* family

Family *Phenuiviridae* categorized in 20 genera, consist of segmented (3 segments) negative sense single stranded RNA viruses infecting plants and animals. Genus *Phlebovirus* associated with disease in ruminants and humans is often vectored by either ticks or mosquitoes. Three novel viruses named Ensuna phenuivirus, Emibwa phenuivirus, and Anopheles phasvirus were identified in Culex spp (*Culex duttoni* and *Culex neavei*), *Culex antennatus*, and *Anopheles coustani* respectively from Arua and Kasese districts. A novel virus named Ensuna phenuivirus was identified in mosquito pools (MosP36 and MosP38) of *Culex duttoni* and *Culex neavei* from Arua district. It was most closely related to Gouleako virus (NC_043051.1) from West Africa; diverging by 24% nucleotide sequence (Marklewitz, 2011 #342). We also detected another novel virus (Emibwa virus) in *Culex antennatus* mosquitoes in pool MosP76 from Kasese district. Emibwa phenuivirus is phylogenetically related to Badu virus (NC_038257.1) first detected in *Culex* spp from Australia (Hobson-Peters, 2016 #343). Nucleotide pairwise distance between Emibwa phenuivirus and Badu virus was 31.61% as shown in the figure below. Anopheles phasvirus another novel virus closely related to Phasi Charoen-like virus (MN866220.1) with 44.29% pairwise distance was detected in pool 5 of *Anopheles coustani* from Arua district. This was distantly related to the Phasi Charoen-like viruses previously detected in *Aedes aegypti* mosquitoes in Karnataka, India and *Aedes aegypti* mosquitoes in Brazil (Aguiar et al., 2016 (Munivenkatappa, 2021 #341).

## Discussion

Uganda is a hotspot for viral pathogens and a site of high viral diversity. In this study, we employed next generation sequencing to highlight the extent of viral richness in Uganda and to identify pathogens of medical and veterinary importance. The last decade has witnessed the identification of several emerging viruses using this method, which could not be identified using traditional PCR- based or culture-based methods, including Ntwetwe virus, Nyangole virus and Kibale virus all from the family *Peribunyaviridae*. This approach is therefore highly likely to lead to the identification of previously uncharacterised viruses, some of which are likely to have the potential to cause disease in human and animal populations. Despite an increase in studies, virus species present in mosquito populations in Sub-Saharan Africa are largely undocumented. This study aimed to investigate the mosquito virome in two geographically distinct areas of Western and North Western Uganda. Virus species diversity was significantly different in the two regions with a richer mosquito virome registered in North western Uganda. A total of 173 viruses in all sampled mosquito genera from multiple virus families including the *Chuviridae, Flaviviridae*, *Togaviridae*, *Rhabdoviridae*, *Peribunyaviridae*, *Phenuiviridae*, *Nairoviridae* and *Orthomyxoviridae*, *Iflaviridae*, *Phasmaviridae*, *Nodaviridae and Alphatetraviridae*, *Iridoviridae*, *Mesoniviridae*, *Nudiviridae*, *Parvoviridae, Permutotetraviridae, Picornaviralsidae, Retroviridae* and *Xinmoviridae* were detected. Out of these, viruses were identified in families known to infect human and animals including *Flaviviridae*, *Orthomyxoviridae, Phenuviridae, Rhabdoviridae, Togaviridae*. Other viruses identified during the study clustered with mosquito borne viruses already described in other parts of the world including Europe, Asia and America. The majority of the sequence contigs detected represent novel species, with unknown potential to cause disease.

This study, carried out in Western and North West Uganda illustrates the scale of richness and virus diversity in the region and the need to further characterise the virome in mosquitoes, especially those with a propensity to feed from human and animal hosts. Out of the 13 novel viruses 10 (76.9%) were from families (*Orthomyxovirida*e, *Phenuiviridae*, *Rhabdoviridae*) associated with disease in humans or animals. Majority (9/13) of the novel viruses were detected in *Culex* species which are previously known vectors of a number of viruses of known medical importance.

Novel viruses identified in family *Phenuiviridae* included Ensuna phenuivirus from Culex species (pool MosP36 and MosP38), Emibwa phenuivirus from *Culex antennatus* (pool MosP76), Anopheles phasivirus from *Anopheles coustani* (pool MosP5). Those identified in family *Orthomyxoviridae* included Anopheles orthomyxo-like virus from *Anopheles spp* (pool MosP61) from Kasese district, Coquillettidia orthomyxo-like virus from *Coquillettidia species* (pool MosP11) in Arua district, Emibwa orthomyxo-like virus from Culex and *Anopheles spp* (pool MosP63, MosP85 AND MosP91) and Mansonia orthomyxo-like virus from *Mansonia uniformis* (pool MosP51 and MosP95) in Arua and Kasese districts. To our knowledge this is the first report of a large-scale study in which NGS is used to characterize viruses from diverse mosquito species in Uganda.

### Limitations to the study

There were several limitations to this study. While we detected near full genomes of viruses in many virus families, metagenomic sequencing can lack sensitivity which can be increased by employing a semi-agnostic target enrichment approach to increase the detection and documentation of novel species. Further studies employing such an approach are indicated. Further, we sampled from two regions of Uganda only. While these areas have varied ecological characteristics (Arua district has an arid climate with lower rainfall while Kasese district has a milder climate with increased rainfall), further sampling in Uganda and indeed in the region is indicated. Finally, while the majority of the viruses described are likely to replicate in mosquitoes and some may be arboviruses, we did not demonstrate vector competence in this study.

---

**Table 1:**
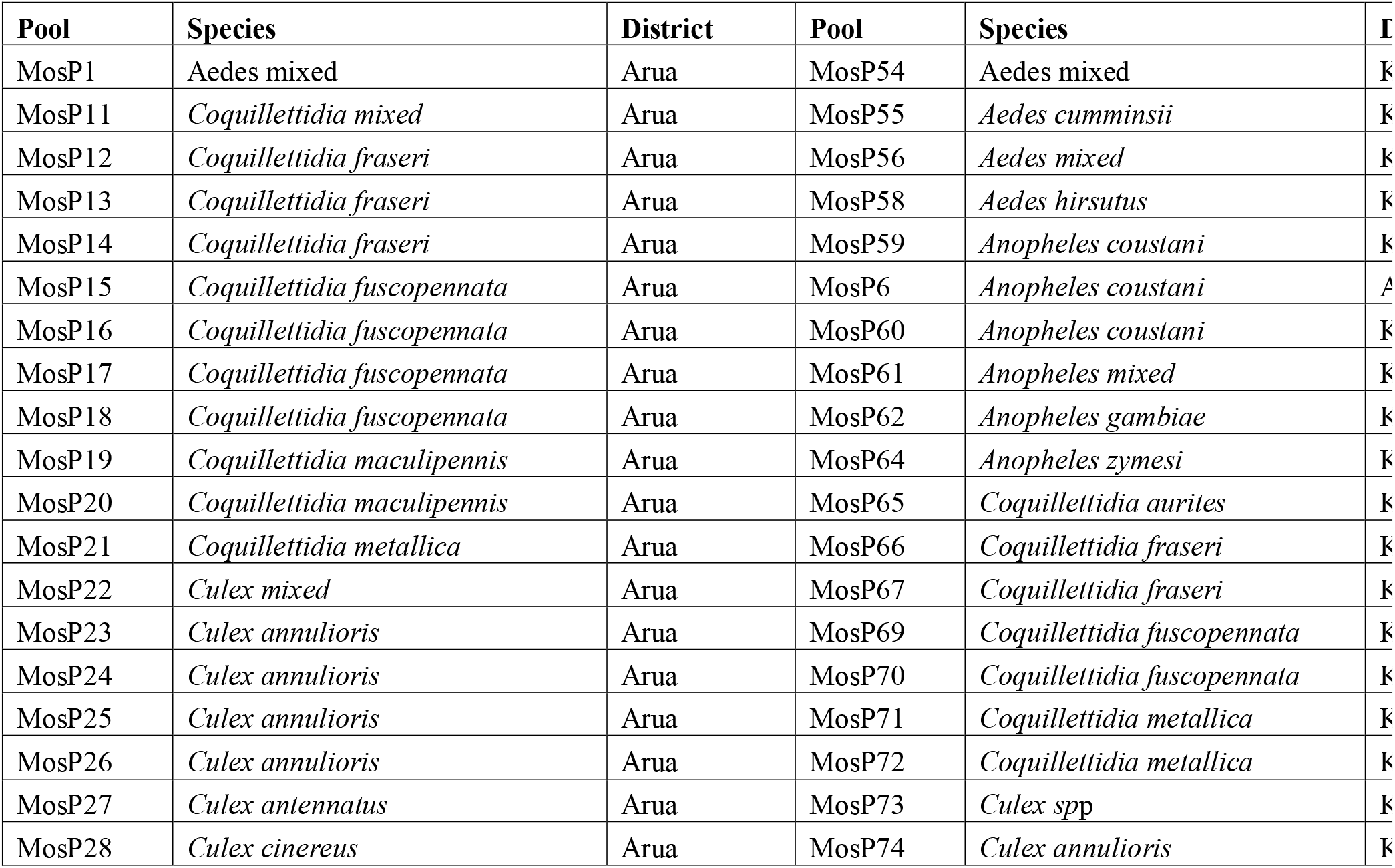
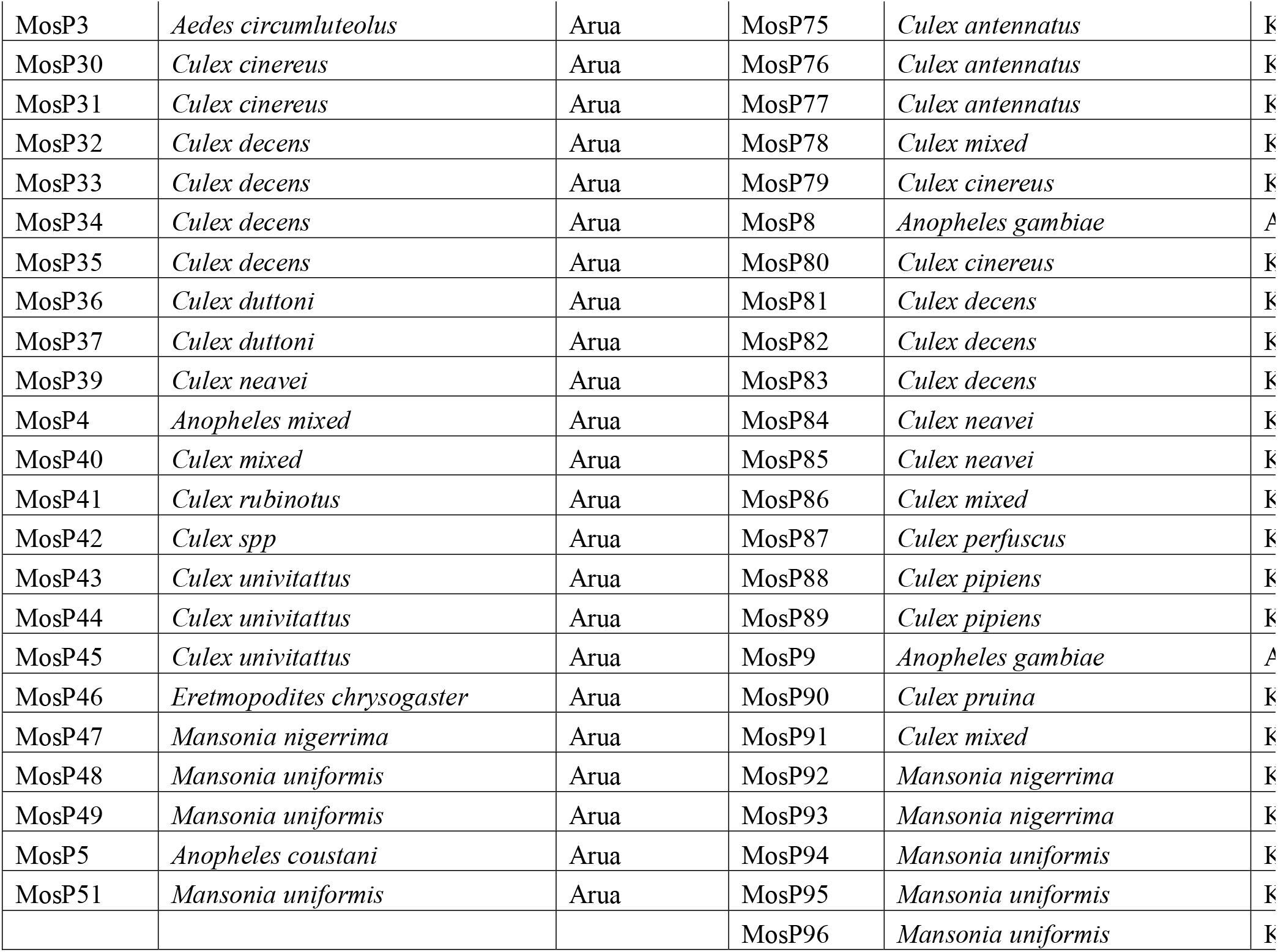
Mosquito pools containing described and unclassified viruses.

**Supplementary Table 1.**
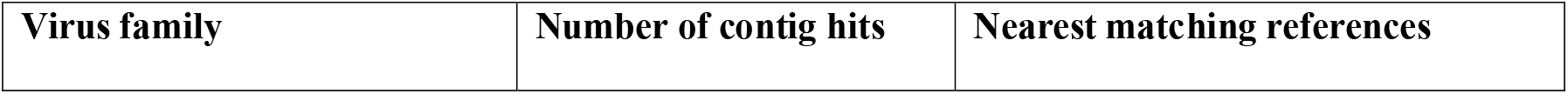
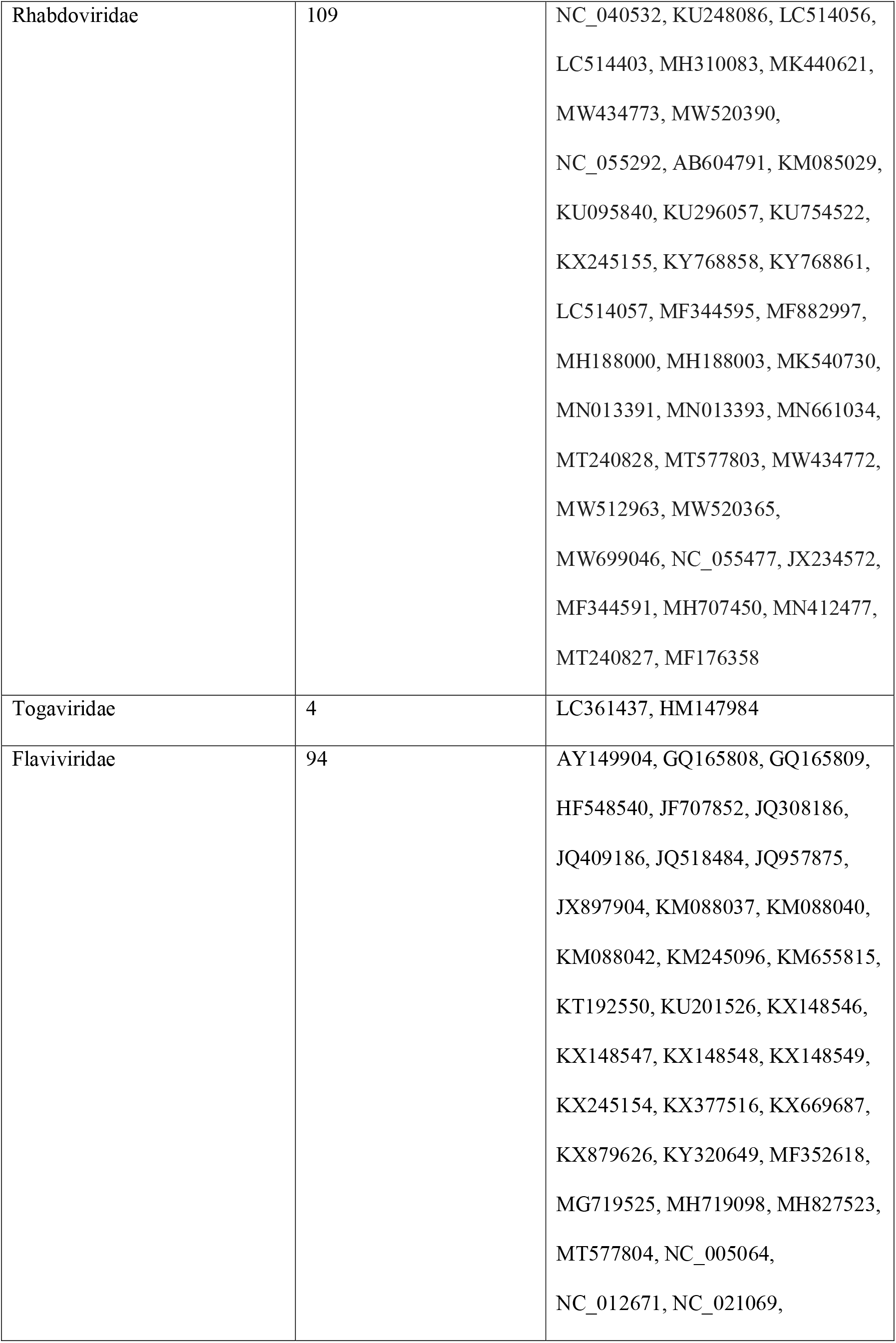
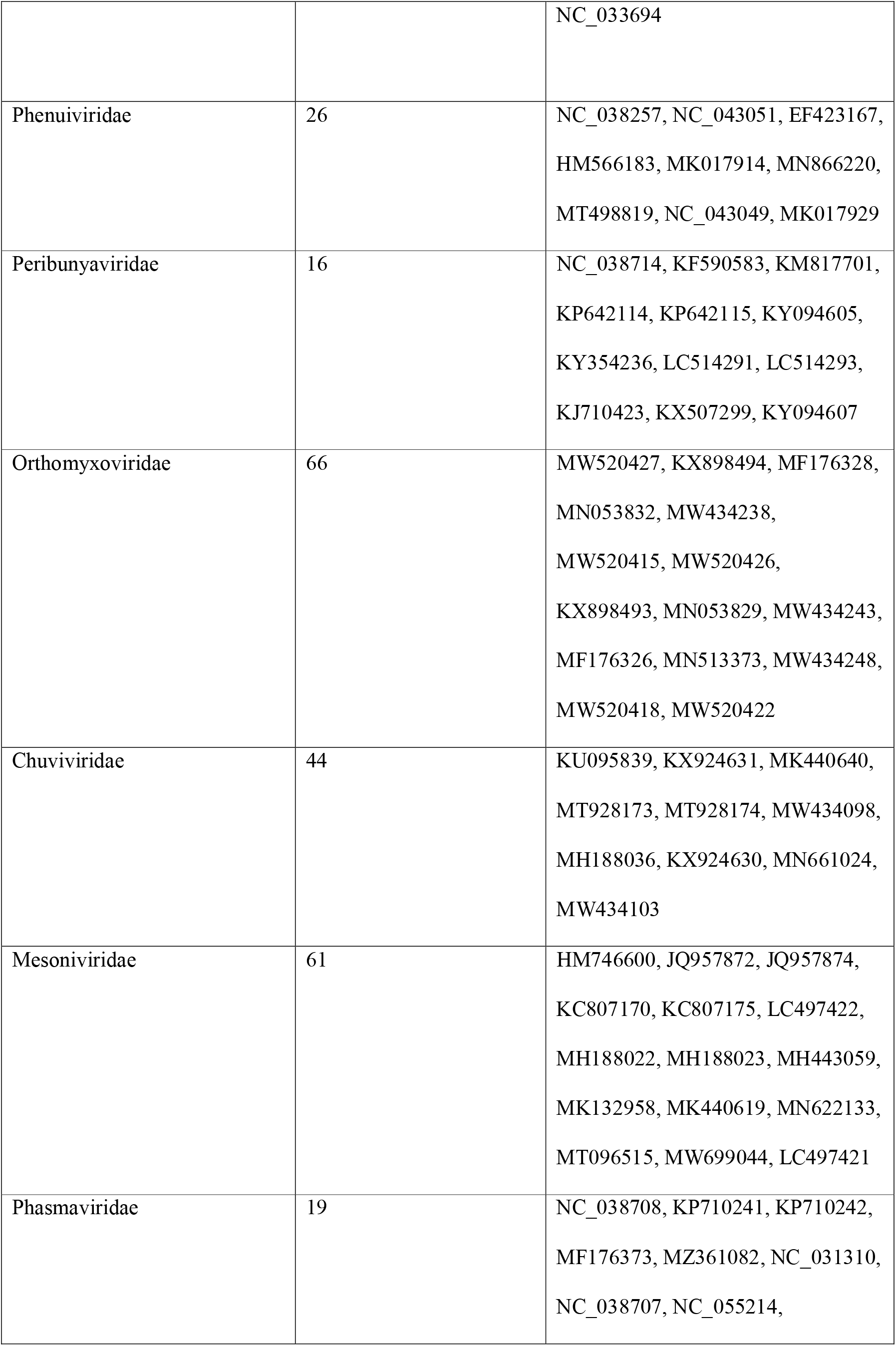
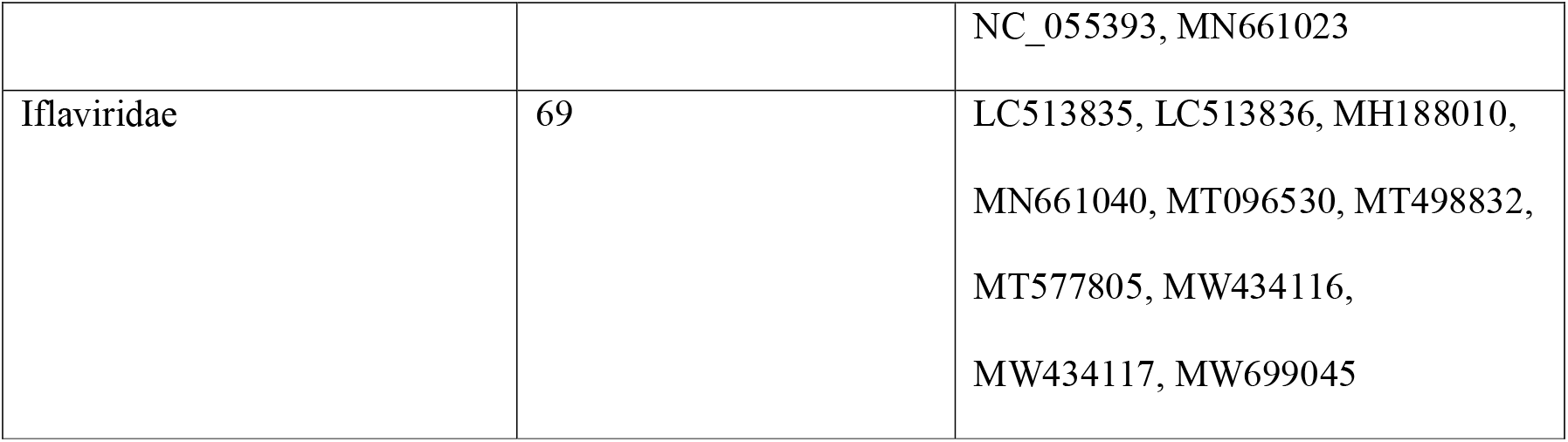

